# Optimization of fermentation conditions for *Metarhizium robertsii* and its effect on rice growth

**DOI:** 10.1101/2025.11.19.689318

**Authors:** Deling Wang, Xiaoyuan She, Meng Zhu, Dongfei Han, Jiaolong Fu, Jing Bai

## Abstract

*Metarhizium robertsii* is an important entomopathogenic fungus and plant endophyte, possessing dual functionality as both an insecticide and a plant growth promoter, which has led to its widespread application in agriculture. However, the effects and underlying mechanisms of *M. robertsii* (MAA)in promoting rice growth remain poorly understood. This study confirmed the growth-promoting effects of MAA and its fermentation supernatant on rice. Based on single-factor and response surface methodology experiments, the fermentation conditions of MAA were subsequently optimized using rice root length as a key indicator, and the potential mechanisms behind this growth promotion were preliminarily investigated. The results demonstrated that both MAA and its fermentation supernatant exerted significant growth-promoting effects on rice. Sucrose and peptone were identified as the optimal carbon and nitrogen sources, respectively. The most pronounced effect was observed when the fermentation supernatant, produced under the conditions of pH 6.0, 0.2% β-Ala, and 100 μmol/L Al_2_(SO_4_)_3_, was applied, resulting in a 21.7% increase in rice root length compared to the control group. Furthermore, significant enhancements in chlorophyll content and denser root hairs were recorded in rice treated with the MAA fermentation supernatant, with the chlorophyll content increasing by 28.17% compared to the control. These findings provide a theoretical basis for the development and application of green and efficient bio-inoculants.

**IMPORTANCE:** *Metarhizium robertsii* is an endogenous plant fungus that can be effective for efficient insect control, but the plant-promoting effect of this fungus and its related studies are less reported. The colonization ability and growth promotion effect of *Metarhizium robertsii* on different crops vary. In this study, a sterile hydroponic system for rice was established. The research found that *Metarhizium robertsii* can colonize the root system of rice, significantly promoting rice growth, and its growtion-promoting effect is affected by the differences in its fermentation products. After the optimization of the response surface of the fermentation conditions of *Metarhizium robertsii*, its growth-promoting ability increased by 21.7%. Developing microbial biofertilizers/biopesticides as a basis for reducing chemical inputs in rice cultivation.

## INTRODUCTION

*Metarhizium robertsii*, as an important class of biocontrol fungi which can host a wide variety of insects, is widely used in agriculture. According to incomplete statistics, the world’s more than 200 kinds of insects can be infected by this fungus to death. With the deepening of research, it has been found that in addition to infecting and destroying target pests, *M. robertsii* can also colonize plant roots and act as a plant endophyte to promote plant growth(1) (2). Therefore, as a microbial resource with a wide range of application prospects, the role of *M. robertsii* in promoting the growth and development of plants has attracted more and more attention.

*M. robertsii* is a symbiont of agriculturally and economically important plants such as tomato, Arabidopsis, soybean, maize and wheat (3-6). When *M. robertsii* colonizes plants, it can provide benefits such as growth promotion (7-9), nutrient transfer (10) and protection against pests (8, 9) and diseases (11, 12). For example, tomato colonized by *M. robertsii* has significantly increased root length and height compared with uncolonized control (13). Arabidopsis fresh weight and total chlorophyll content significantly increased 7 d post-inoculation with the three *Metarhizium anisopliae* strains Ma-20, Ma-25 and Ma-28 evaluated. The primary root length was promoted, and β-caryophyllene and o-cymene were identified as metabolites during this interaction. However, the research on the plant growth promoting effect of *M. robertsii* is still immature, and the scope of researched plants is limited. Especially, there are only a few reports on the interaction between *M. robertsii* and rice, and they focus on the stress resistance of rice to heavy metal and salt (14-16). The endophytic and growth promoting effects of *M. robertsii* on rice remain unclear.

The production of secondary metabolites is the key to plant growth promoting effect. For example, Pipecolic acid (PIP) could form a non-selective rhizosphere relationship between *M. robertsii* with Arabidopsis Thaliana. The PIP level of Arabidopsis increased after the inoculation with *M. robertsii*, which enhanced the resistance to plant pathogens and aphids (17). Destruxins (DTXs), including 25 DTXs analogues, were detected under the conditions of co-culture of 4 *M. robertsii* with soybean and corn. At the same time, it was found that there were significant differences in DTXs yield under different plant conditions (18). Kumar showed that Metarhizium can produce a range of organic acids, such as keto-gluconic, oxalic, which promotes Zinc solubilization, and a range of secondary metabolites such as IAA iron carriers, which promote the growth of rice and cardamom plants (19).

In addition, the output of these bioactive secondary metabolites is not only related to the genetic characteristics of the fungus itself, but also to the culture medium, nutrient composition, and fermentation conditions(20). Optimizing the physical and chemical environmental factors that affect a fermentation process (such as temperature, pH, aeration, and medium composition) is crucial for maximizing the yield of a specific bioproduct (21). By optimizing the composition of the medium and the fermentation conditions, secondary metabolites with growth-promoting effects can be significantly increased (22). He et al. showed that optimal fermentation conditions of *M. robertsii* increased the inhibition rate of *Fusarium solani*, which is the main pathogenic fungus causing the root rot of wolfberry (*Lycium barbarum*)(23). Through response surface methodology and metabolomics, a maximum indole auxin production of 208.3 ± 0.4 mg IAA equ/L by *Pantoea agglomerans* was obtained, which was a 40% increase compared to the growth conditions used in previous studies (24). Response Surface Methodology (RSM) has been widely exploited to optimize fermentation processes, but surprisingly it has been rarely used in combination with metabolomics, particularly untargeted metabolomics. Furthermore, few studies are available on the process optimization of secondary metabolites production in *M. robertsii* for plant growth-promotion.

The objectives of this study were to (1) analyze the effects of different carbon and nitrogen sources and fermentation conditions on the growth-promoting effect of *M. robertsii*; (2) The response surface method was used to optimize the fermentation conditions of *M. robertsii* to improve the growth-promoting effect of the fermentation supernatant; (3) To explore the possible mechanism of growth promotion on rice.

## RESULTS

### Effects of MAA Fermentation Supernatant on Rice Growth Promotion

The growth-promoting effects of MAA fermentation supernatant were evaluated using a hydroponic rice seedling system. As shown in Figure 1, comparative analysis between the treatment group and the non-inoculated control group revealed significant morphological differences after 7 days of cultivation. Seedlings treated with MAA fermentation supernatant showed a 13.55% increase in shoot elongation compared to control (P < 0.05), reaching an average height of 16.08 cm versus 14.16 cm in control. Simultaneously, primary roots exhibited 1.164 times greater elongation and adventitious root formation was enhanced by 12.5% compared to control (P < 0.05).

**Figure 1.**
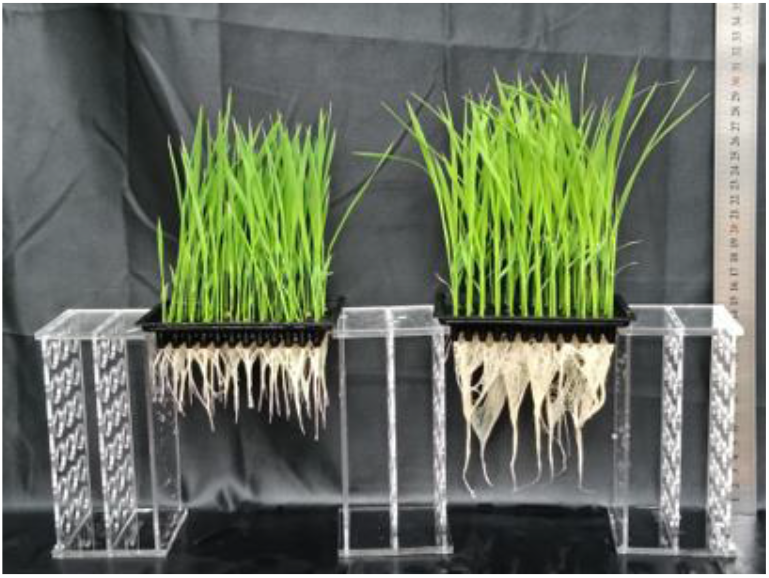
Growth-promoting effect of on Rice (left) control group (right) test group.

### Effect of MAA on the Abundance of Endophytic Fungi in Rice Roots

Following the observed growth-promoting effects of MAA on rice, we further investigated whether MAA could colonize rice roots and modulate their endophytic fungal communities. Using ITS region amplification and high-throughput sequencing, we systematically analyzed the impact of MAA inoculation on the root-associated fungal microbiome. The results showed that the MAA treatment group (M^+^) showed a 400-fold increase in MAA relative abundance compared to the control group (M^-^) (from 0.00045% to 0.18%; p < 0.01). This represented a 99.75% increase in detection frequency (Figure 2). As shown in Figure 2, the M^+^ group significantly altered the composition of the endophytic fungal community. Notably, the relative abundance of *Fusarium* sp.—a pathogenic genus associated with root rot—was markedly reduced by 76.15% (p < 0.05). Similarly, *Dictyostelium* sp. (chlorosis-associated) and *Pezizaceae* sp. (associated with chlorotic lesions) declined by 47.81% and 89.67%, respectively. Additionally, the MAA treatment inhibited the proliferation of *Gibberella* sp., a soilborne pathogen responsible for rice seedling blight. Conversely, the treated roots exhibited a significant increase in beneficial fungal genera: *Trichoderma* sp. (a plant growth-promoting taxon) and *Byssochlamys* sp. (known to suppress fungal pathogens) were enriched by 79.71% and 0.0032%, respectively (p < 0.01) (Figure 2). These findings suggest that MAA successfully colonizes the rice rhizosphere and enhances rice growth by restructuring the root-associated fungal community, enriching beneficial symbiotic and antagonistic taxa while significantly suppressing pathogenic fungi in the root endophytic community.

**Figure 2.**
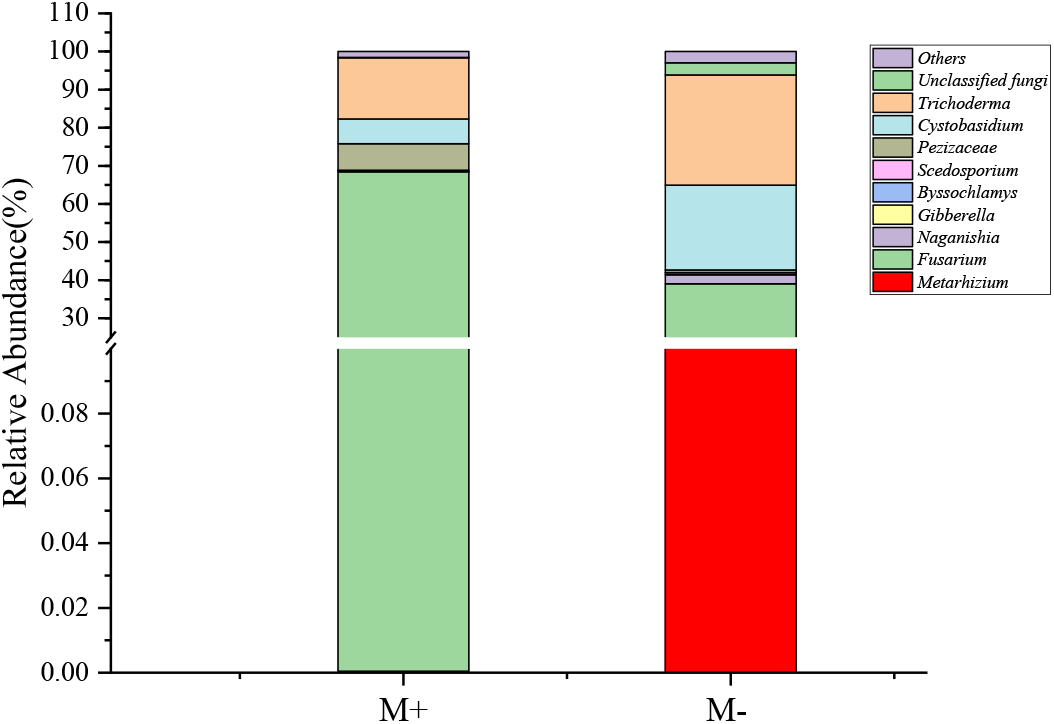
Relative abundance of dominant genera in root samples from different treatment. Note: M+: rice root inoculated with MAA in rhizosphere soil; M-: rice root without treatment.

### Effects of Culture conditions on MAA growth and Rice Growth Promotion

To systematically evaluate the effects of culture conditions on MAA growth and its rice growth-promoting potential, we selected six critical variables (carbon source, nitrogen source, pH, cultivation time, β-Ala concentration, Al_2_(SO_4_)_3_ supplementation) for optimization, which could critically influence fungal growth and metabolite production. Specifically, β-Alanine (β-Ala) may enhance the secretion of phytohormones and siderophores, influencing metabolic pathways in fungi(25). In plant-associated microbes, β-Ala derivatives have been implicated in stress tolerance and growth promotion. Aluminum sulfate (Al_2_(SO_4_) _3_) acts synergistically to modulate metal ion availability, potentially affecting fungal metabolite secretion and nutrient acquisition. Using the basic fermentation medium as a control, the supplementation of sucrose and peptone as carbon and nitrogen sources increased plant height and root length by 24.59%, 22.55%, and 77.17%, 72.73%, respectively, which was superior to other treatment groups (p<0.05) (Figure 3A and 3B). In addition, compared with the control group, supplementing 0.2% β-Ala resulted in an increase of 28.47% and 76.64% in plant height and root length, respectively (p<0.01). The addition of 50 μmol/L Al_2_(SO_4_)_3_ increased plant height and root length by 27.51% and 68.04%, respectively (Figure 3B). After 7 days of cultivation in the fermentation broth, there were significant differences in plant height and root length between the hydroponic group and the control group, with increases of 29.97% and 71.19%, respectively. Furthermore, when the pH was adjusted to 6, plant height and root length increased by 29.86% and 75.54% compared to the base medium. (Figure 3B). These results collectively demonstrate significant variations in the plant-growth-promoting effects of MAA fermentation supernatant under different culture conditions.

**Figure 3.**
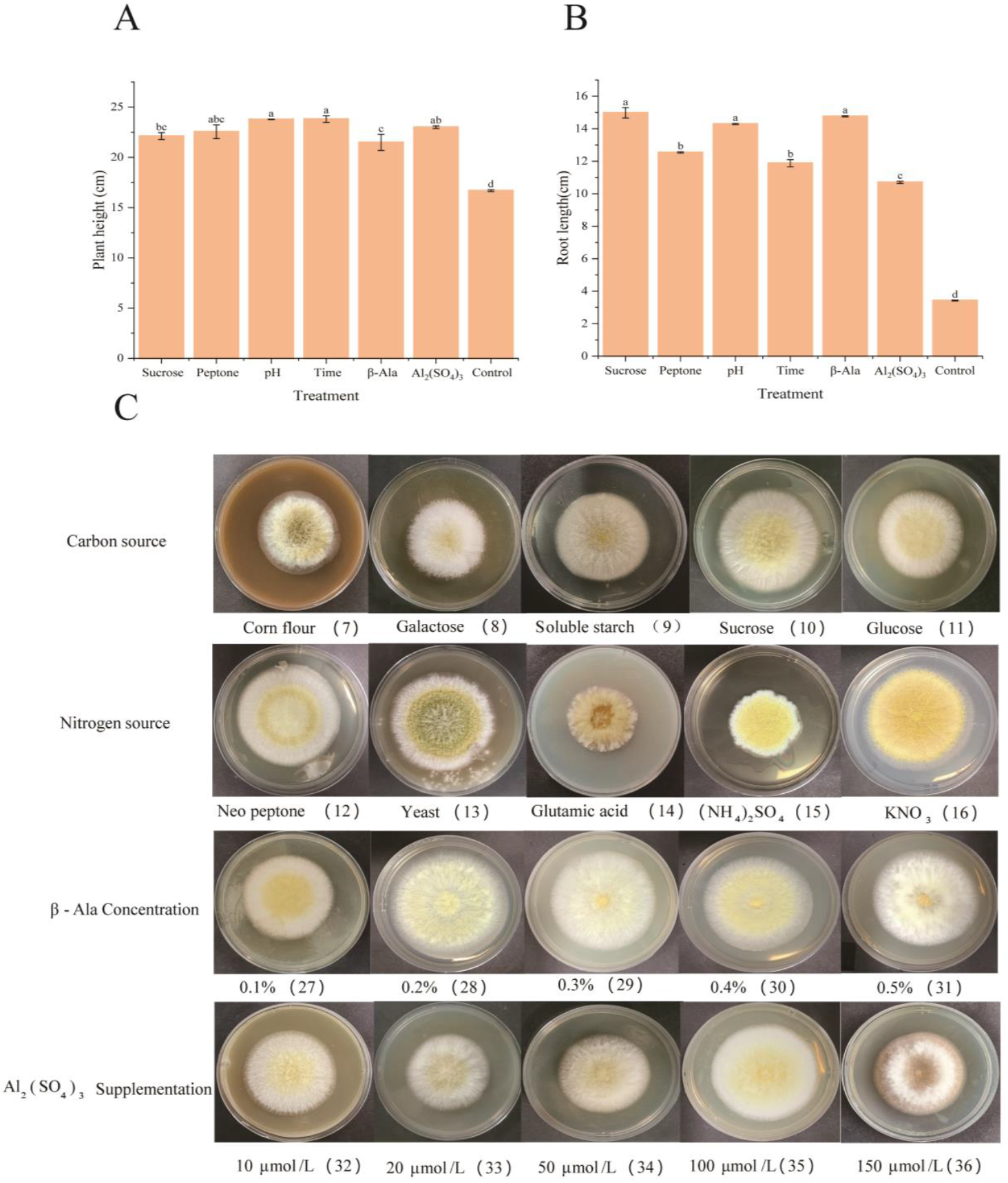
Effects of different culture conditions on Rice seedling growth and the growth of MAA. (A) and (B) respectively represent the effects of MAA fermentation culture in different liquid media on the plant height and root length of Rice. Control indicates the culture of MAA in the basal medium, and treatment indicates the optimal value under the single-factor experimental design. The horizontal coordinates sucrose and peptone respectively represent the replacement of carbon and nitrogen sources in the basic culture medium, pH indicates the alteration of the pH value of the culture medium, time represents the change of the fermentation time of MAA, and β-Ala and Al_2_(SO_4_)_3_ respectively represent the addition of different concentrations of β-Ala and Al_2_(SO_4_)_3_; (C) Effect of different culture methods on solid medium. The influence of different solid media on the growth of MAA. The numbers in parentheses indicate the numbering of culture media in Table S4.

**Figure 4.**
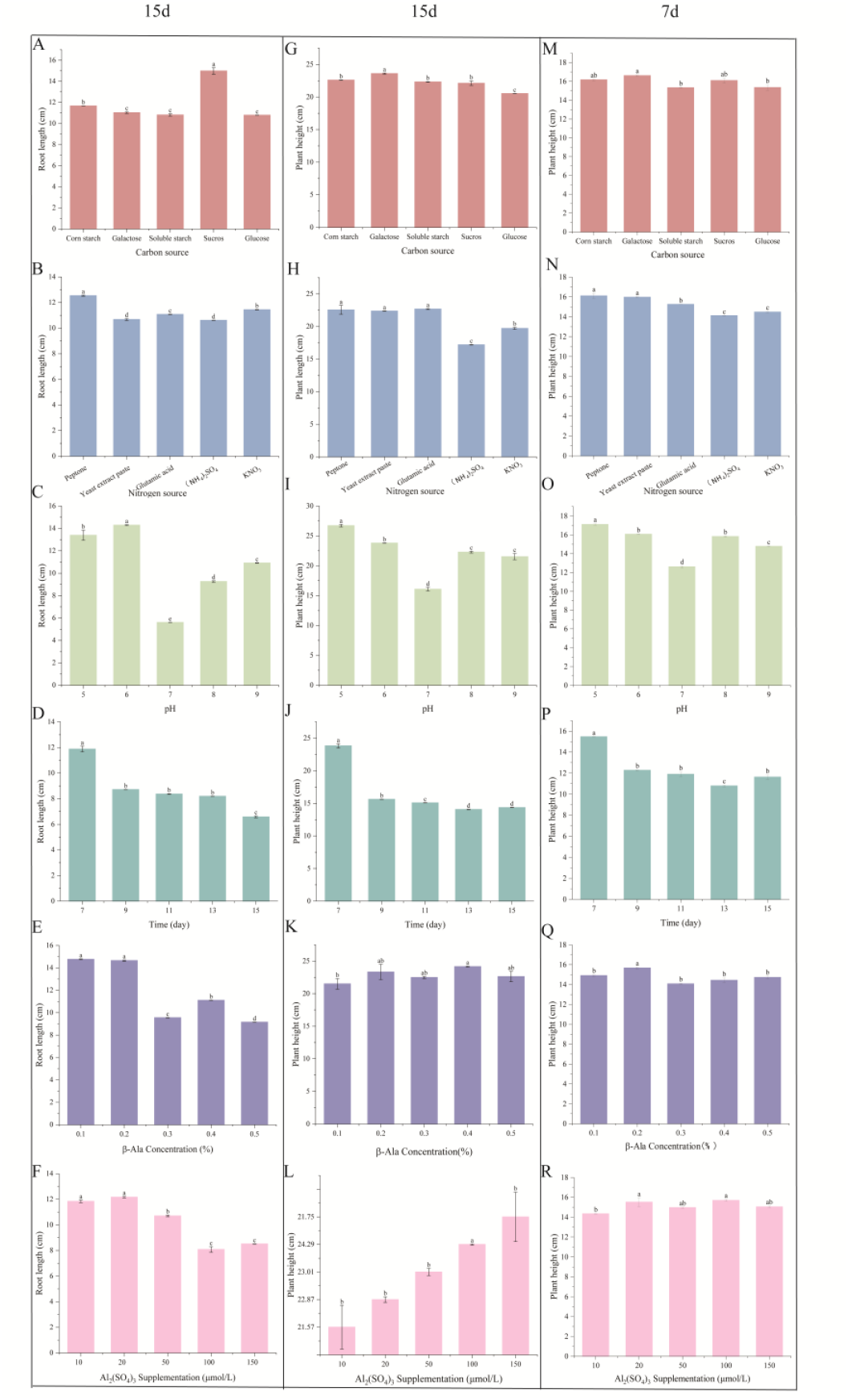
Growth responses of Rice seedlings to *M. robertsii* fermentation variables. (A) Fermentation time, (B) β-Ala concentration, (C) Al_2_(SO_4_) _3_ concentration, (D) carbon source, (E) nitrogen source, and (F) pH on plant height (7/15 d) and root length (15 d). Error bars: ±SE; different letters denote significant differences (p < 0.05).

Observing the effect of different culture medium components on the growth of MAA strain through colony morphology. The corresponding media compositions are listed in Appendix Table S4. colony morphology exhibited significant variation across culture conditions (p <0.05), with diameter measurements demonstrating clear media-dependent growth patterns (Figure. 3C). The numbers in parentheses indicate the numbering of culture media in the appendix list Table S4. The carbon source screening results showed that sucrose (10) supported maximum growth (diameter 15.82 cm), significantly better than other carbon sources. The colonies were circular with neat edges, and the surface of the colony is dry, presenting a dense fuzzy texture. Peptone (12) produced extensive colonization (diameter 15.50 cm) and a unique white mycelial morphology, with mature fungal colonies covered in a large number of spores in a distinct powder form. The medium with 0.2% β-Ala produced the largest colonies (28) (17.73 cm), and the hyphae showed obvious spreading growth. The Al_2_(SO_4_)_3_ tolerance test showed that when 100 μmol/L (35) was added, the colony size reached its maximum of 16.11 cm (Figure 3C). For detailed information and descriptions, refer to Supplementary Material Table S5. Systematic evaluation using standardized media formulations (Table S4) revealed significant culture-dependent variations in both MAA colony growth and rice growth promotion, and a positive correlation was observed between the extent of MAA growth stimulation and the efficacy in promoting rice growth. Based on these findings, key parameters including carbon source, nitrogen source, pH, fermentation duration, β-Ala concentration and Al_2_(SO_4_)_3_ supplementation were selected for single-factor optimization experiments to enhance the plant growth-promoting capacity of MAA.

### Single-Factor Test

The fermentation broth’s efficacy exhibited distinct response patterns to individual culture parameters. Fermentation time exhibited an inverse relationship with growth promotion. The 7-day treatment yielded the optimal plant height of 15.46 cm. In contrast, extended incubation to 15 days shifted biomass allocation toward shoot growth (23.82 cm) at the expense of root development (11.88 cm). β-Alanine concentration followed a biphasic trend, peaking at 0.2% with plant heights of 15.66 cm (7 d) and 14.65 cm (15 d roots)—representing 12.3% and 23.1% increases versus 0.1%, respectively (p < 0.05). Aluminum sulfate displayed concentration-specific regulation, where 100 μmol/L maximized plant height (15.69 cm at 7 d; 24.29 cm at 15 d), while 20 μmol/L optimized root development (11.9 cm). Nutrient sources differentially modulated growth: sucrose yielded maximum 15-day root elongation (14.98 cm), whereas peptone enhanced both plant height and root metrics (16.11 cm 7-d height; 12.54 cm 15-d roots). pH effects culminated at pH 6 with 23.79 cm root length, demonstrating narrow optimal ranges for metabolic activity. All treatments maintained biological reproducibility (SEM ±0.15-0.45 cm, n=3), with non-linear responses underscoring the precision required in fermentation parameterization.

### Response Surface Test Optimization Results

The response surface test’s design and results are shown in Table S2.

### Regression Equation Fitting and Analysis of Variance

A regression model for root length was established based on statistical analysis of the experimental data, as expressed in the following equation: Root length = 12.14 + 0.2739 × A + 0.0044 × B - 0.1472 × C - 0.0756 × D + 0.0875 × AB + 0.1888 × AC - 0.1000 × AD + 0.2038 × BC - 0.0200 × BD + 0.0688 × CD - 0.2243 × A^2^ - 0.6093 × B^2^ - 0.6643 × C^2^ - 0.5793 × D^2^.

Variance analysis and a significant difference test were conducted for the regression model, and the results are shown in Table S3. The model F-value of 16.11 implies that the model is significant. There is only a 0.01% chance that an F-value this large could occur due to noise. P-values less than 0.0500 indicate that model terms are significant. In this case, A, AC, BC, B^2^, C^2^, and D^2^ are significant model terms. Values greater than 0.1000 indicate that the model terms are not significant. If there are many insignificant model terms (not counting those required to support hierarchy), model reduction may improve the model. The lack-of-fit F-value of 1.94 implies that the lack of fit is not significant relative to the pure error. There is a 24.12% chance that a lack-of-fit F-value this large could occur due to noise. A non-significant lack of fit is good—we want the model to fit. The equation in terms of coded factors can be used to make predictions about the response for given levels of each factor. By default, the high levels of the factors are coded as +1 and the low levels are coded as −1. The coded equation is useful for identifying the relative impact of the factors by comparing the factor coefficients.

### Response Surface Analysis of Interaction of Various Factors

The response surface diagrams intuitively reflect the interactions between the key factors and their influence on rice root length. From the surface plots and contours in Figure 5, the interactions between sucrose concentration (A), pH (B), β-Ala concentration (C), and Al_2_(SO_4_)_3_ supplementation (D) and their significant effects on root length can be observed. This observation is consistent with the ANOVA results shown in Table S3, where interaction terms AC (sucrose concentration × β-Ala) and BC (pH × β-Ala) were identified as significant model terms (p < 0.05).

**Figure 5.**
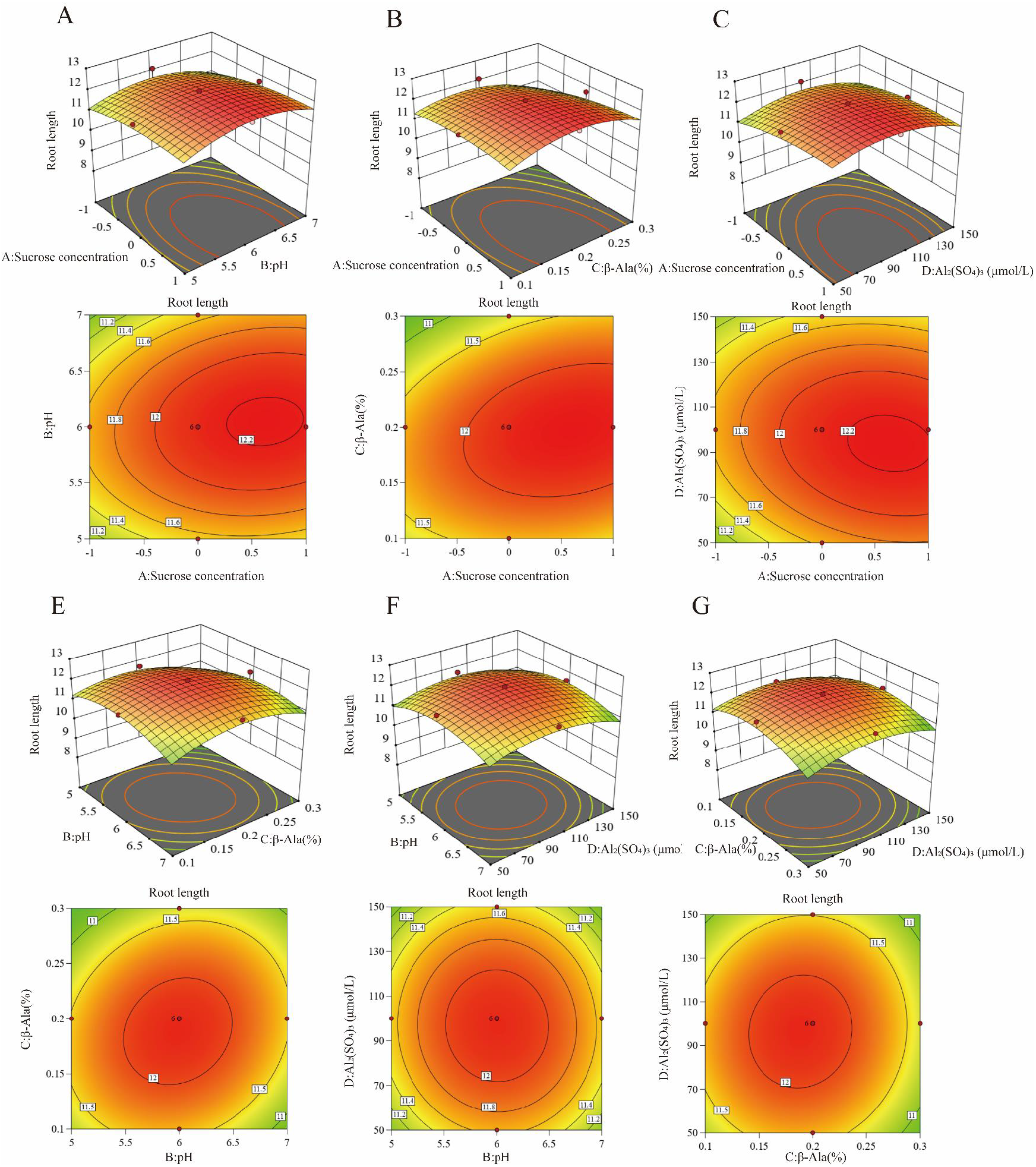
The response surface methodology and contour plots of the effects of the interaction between pH and sucrose concentration (A), β-Ala and pH (B), β-Ala (C) and pH, pH and β-Ala, pH and Al_2_(SO_4_)_3_, sucrose concentration and Al_2_(SO_4_)_3_ supplementation (D) on the root length of the strain fermentation broth. Note: The incline degree of the surface diagram is directly proportional to the influence degree of factors on the response value. The larger the focal length of the contour, the stronger the interaction between parameters.

The optimal conditions were obtained using a quadratic polynomial regression fitting equation: sucrose concentration = 26 g/L; pH = 6.03; β-Ala concentration = 0.20%; and Al_2_(SO_4_)_3_ supplementation = 96.07 μmol/L. Under these conditions, the predicted root length was 12.23 cm. For simple and feasible operation, the optimal fermentation conditions were determined as follows: sucrose concentration of 30 g/L, pH of 6.0, β-Ala concentration of 0.2%, and Al_2_(SO_4_)_3_ concentration of 100 μmol/L.

### Establishment and Validation of Optimal Culture Conditions

The model regression equation was resolved using the software to derive the optimum culture conditions: The fermentation broth was prepared under the optimal fermentation conditions sucrose concentration was 26 g/L, pH 6.0, β-Ala 0.2% and Al_2_(SO_4_)_3_ 100 μmol/L and repeated three times to verify the accuracy of the model. The average root length was 11.95 cm, which was consistent with the predicted value of 12.23 cm.

### Effect of Different Cultures on Root Hairs and SPAD in Rice Roots

In rice, root hairs constitute an essential part of the root system, contributing to nutrient and water uptake and transport, as well as plant anchorage (26). These functions underscore the critical role of root hairs in rice growth and development. SPAD values, which serve as an indicator of chlorophyll content, are widely used to assess photosynthetic efficiency in green plants (27). They reflect key physiological processes including plant growth and development, metabolic changes, and nutritional status, and are commonly employed as a reference parameter in environmental physiology studies. Consequently, chlorophyll content is frequently adopted as a physiological marker to investigate plant growth and development.

Root hair morphology observations revealed distinct differences between the untreated control (Figure. 6A) and the optimal MAA treatment group (Figure. 6B) (selected based on an integrated assessment of all physiological parameters). As demonstrated in Figure. 6, plants treated with MAA exhibited significantly denser and longer root hairs compared to the blank control, indicating enhanced root system development and improved capacity for nutrient and water acquisition. SPAD values in rice plants treated with MAA fermentation broth initially decreased and then increased with prolonged fermentation time, showing the most pronounced effect and the highest chlorophyll reading (29.723) at 7 days of fermentation compared to the blank control (Figure. 7A). During the treatment period, SPAD values exhibited a trend of initial increase followed by a decrease with rising β-Ala concentration. The peak was observed at 0.2% β-Ala, with chlorophyll contents of 30.513 and 30.3 recorded at 0.2% and 0.4% β-Ala, respectively (p < 0.05) (Figure. 7B). As shown in Figure. 7C, the addition of Al_2_(SO_4_)_3_ at various concentrations significantly enhanced chlorophyll content compared to the control, with the most notable effect at 10 μmol/L (chlorophyll content = 30.94), followed by 100 μmol/L (approximately 28.73). Furthermore, different carbon sources in the MAA culture differentially influenced chlorophyll levels, with corn flour and sucrose yielding the most marked effects (approximately 28.29 and 28.54, respectively). Variations in nitrogen sources also affected chlorophyll content, among which (NH_4_)_2_SO_4_ resulted in the highest value (31.41). Finally, the pH of the fermentation broth significantly modulated chlorophyll accumulation, with pH 7 inducing a substantially greater response (chlorophyll content = 33.33) than other treatments.

**Figure 6.**
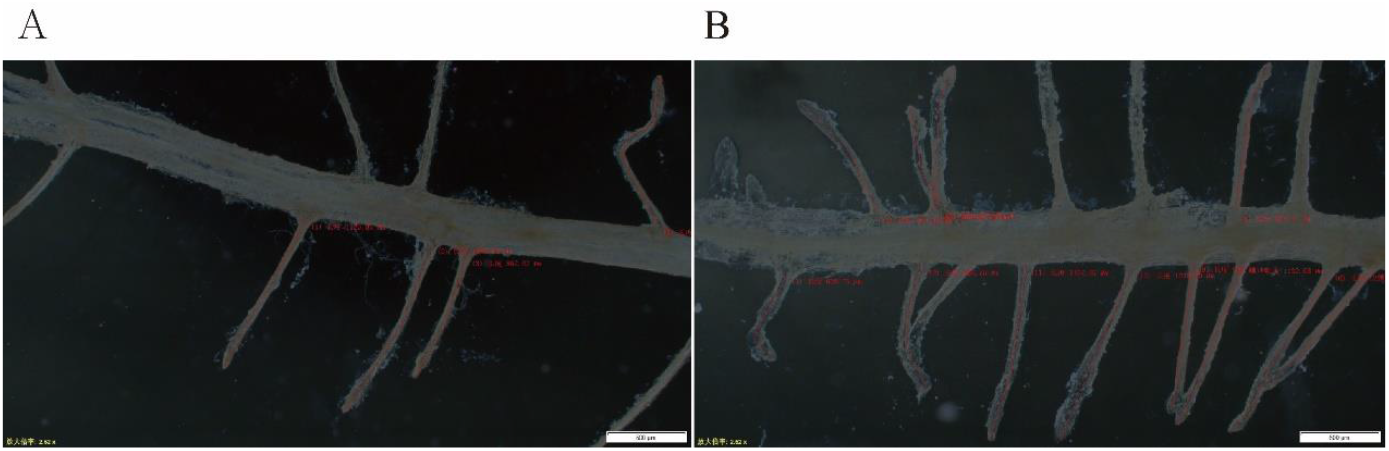
Effects of different MAA fermentation supernatants on rice root hair development. (A) Control: rice treated with supernatant from MAA basal medium after 7-day fermentation. (B) Treatment: rice treated with supernatant from the optimal medium (sucrose, pH 6, 0.2% β-Ala, and 100 μmol/L Al_2_(SO_4_)_3_) after 7-day.

**Figure 7.**
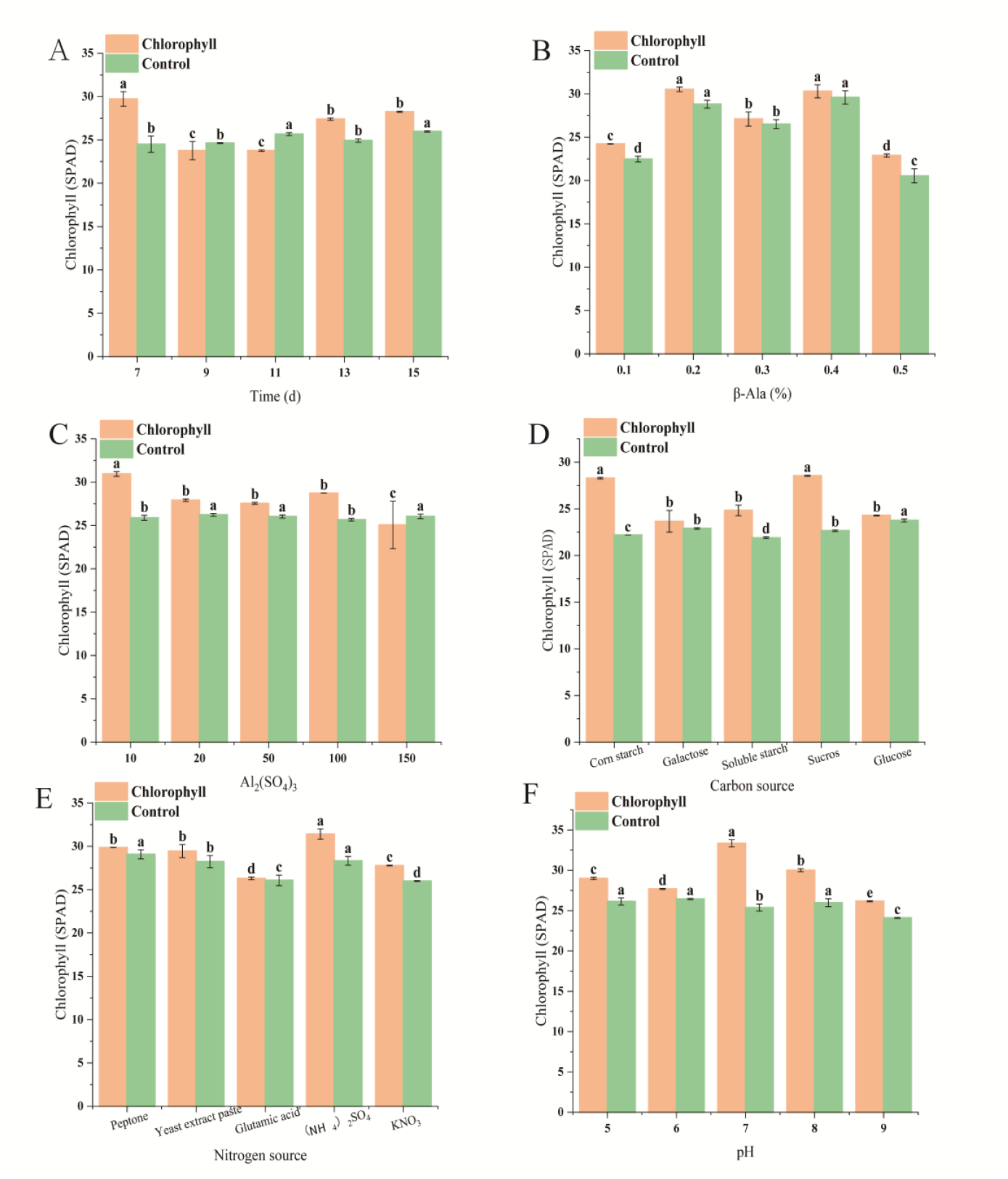
Effects of different cultures of fermentation time(A), β-Ala concentration(B), Al_2_(SO_4_)_3_ supplementation (C), carbon source (D), nitrogen source (E), and pH(F) on SPAD of Rice plants at 15 d.

In summary, these findings indicate that MAA fermentation duration, along with the concentrations of β-Ala and Al_2_(SO_4_)_3_, the type of carbon and nitrogen sources, and the pH, all significantly regulate photosynthetic capacity and root system development in rice, as reflected by SPAD values and root hair morphology. Based on comprehensive evaluation, the optimal conditions for enhancing these key physiological and morphological traits were determined as follows: 7 days of fermentation, 0.2% β-Ala, 100 μmol/L Al_2_(SO_4_)_3_, sucrose as the carbon source, peptone as the nitrogen source, and pH 7.

## DISCUSSION

The direct growth-promoting effects of *Metarhizium robertsii* (MAA) and its fermentation supernatant on rice were first established in this study. Our hydroponic experiments demonstrated that the application of MAA fermentation supernatant significantly enhanced key growth parameters, including root length, plant height, and chlorophyll content. This aligns with the well-documented role of *Metarhizium spp*. as plant root symbionts capable of promoting growth in various crops(7). The observed phenotypic enhancements are likely facilitated by the successful colonization of MAA in the rice rhizosphere, as confirmed by our ITS sequencing analysis which showed a 400-fold increase in MAA abundance in treated roots. This endophytic establishment is a hallmark of *Metarhizium*-plant interactions, forming the foundation for subsequent growth promotion and pathogen suppression(28). The restructuring of the root fungal community, notably the enrichment of beneficial *Trichoderma* and the suppression of pathogenic *Fusarium*, further underscores a potential indirect mechanism through which MAA fosters a conducive growth environment. The optimization of culture conditions for MAA has been shown to significantly enhance its growth and efficacy as a plant growth promoter. In this study, we identified that a sucrose concentration of 26 g/L and a pH of 6.0 were optimal for the growth of MAA, which is consistent with previous findings (29, 30). However, our results also indicated that the addition of specific compounds, such as β-Ala concentration and Al_2_(SO_4_)_3_ supplementation, further improved the MAA’s biomass production and secondary metabolite synthesis. These findings suggest that tailored culture conditions can significantly enhance the bioactivity of MAA, making it a more effective biofertilizer.

The integration of β-Ala and Al_2_(SO_4_)_3_ into the basal medium, adjustments via response surface methodology (RSM), represents a significant advancement in MAA fermentation optimization. The RSM-derived optimal conditions (pH 6.0, 26 g/L sucrose concentration, 0.20% β-Ala, 100 μmol/L Al_2_(SO_4_)_3_) enhanced rice root length to 12.23 cm, increased 21.7% compared to the control. Notably, β-Ala, a precursor of coenzyme A, likely facilitated the synthesis of growth-promoting metabolites such as indole derivatives (31), while Al_2_(SO_4_)_3_ may have acted as a pH stabilizer and elicitor for chitinase production — a hypothesis supported by the 1.7-fold increase in chitinolytic activity (p < 0.01) in the modified medium. These findings align with studies highlighting amino acid supplements as critical drivers of fungal secondary metabolism (32), yet the antagonistic effects of excessive aluminium on sporulation (>0.2 mM) underscore the necessity for precise dosage control.

Hydroponic experiments unequivocally demonstrated the biostimulatory potential of MAA fermentation broth. After 7 days of treatment, the plant height and root length of rice seedlings increased by 35.93% and 44.32%, respectively, compared with the control group; the chlorophyll content increased by 28.29%, and the lateral root formation was abundant (9.6 ± 2.1/plant, compared to 4±1.4 roots/plant in the control group). The successful endophytic colonization of MAA within rice roots, as confirmed by our ITS sequencing results, likely serves as the foundation for these observed growth enhancements. Such phenotypic enhancements may stem from dual mechanisms: (i) fungal-derived auxin analogs directly stimulating cell elongation and root branching, and (ii) improved nutrient solubilization via organic acid secretion, which facilitates phosphorus and iron uptake.

Strikingly, ITS sequencing revealed a 37% reduction in pathogenic Fusarium spp. abundance and a 2.1-fold enrichment of beneficial Trichoderma in the rhizosphere of treated plants. This microbiome restructuring, potentially triggered by MAA-secreted antibiotics (e.g., destruxins) or competition for niche colonization, suggests an indirect pathway for plant growth promotion. However, the observed shift in arbuscular mycorrhizal fungi (AMF) colonization also warrants attention, as AMF play a critical role in phosphorus uptake and accumulation in plants. Future studies should evaluate whether MAA metabolites influence AMF functionality and phosphorus acquisition, and whether they interfere with strigolactone signaling, a key regulator of AMF symbiosis (33).

Despite some progress, constraints persist. Hydroponic systems inherently lack soil-microbe complexity, potentially overestimating field efficacy. Additionally, while β-alanine improved fungal performance, its economic viability for large-scale fermentation requires techno-economic analysis. Further investigations should prioritize: (i) field trials under diverse agroclimatic conditions, (ii) metabolomic profiling to identify bioactive compounds, and (iii) co-inoculation trials with AMF to balance pathogen suppression and nutrient uptake synergies.

## MATERIALS AND METHODS

### Fungal Strain, Culture Medium, and Plant

The fungus *Metarhizium robertsii ARSEF 23* (MAA) was kindly provided by Prof. Yuquan Xu from Biotechnology Research Institute, Chinese Academy of Agricultural Sciences. The strain was cultured on potato dextrose agar (PDA) (BD Difco™) plates at 28°C for 7 d to obtain fresh conidia. Potato dextrose broth (PDB) medium (BD Difco™) was prepared to obtain the MAA fermentation suspension for rice growth promotion determination, and MAA fermentation conditions were optimized using a basic fermentation medium. The composition was as follows: peptone (6 g), K_2_HPO_4_ (0.5 g), glucose (10 g), KH_2_PO_4_ (0.5 g), MgSO_4_ (0.01 g), and distilled water (up to 1 L). Different carbon and nitrogen sources were used to replace glucose and peptone in the basic medium.

*Oryza sativa L*. Shuhui 527 (SH527) seeds, an elite indica hybrid rice restorer in China, were used for our experiments.

### Determination of the Growth promotion Effect of MAA Strain on Rice

Transfer 1 mL of spore suspension of MAA strain (1 × 10^7^ CFU/mL) into 100 mL of PDB medium in 250 mL flasks and incubate the culture at 28 °C, 220 rpm for 7 d. The post-fermentation broth was centrifuged at 4 °C and 8000 rpm for 5 min to obtain the supernatant, which was then sterilized through a 0.22 µm filter and set aside.

Rice seeds (SH527) with full grains and uniform morphology were selected and treated with 75% alcohol and 10% NaClO solution for surface disinfection, and then washed with sterile water three times. The disinfected seeds were immersed and maintained in a constant temperature incubator at 28 °C for 8 h, then transferred to a petri dish covered with sterile filter paper and kept in the dark for 3 d, and seedlings with consistent germination status (coleoptile length 1.0 ± 0.2 cm) were screened for use. Black polyethylene hydroponic box (specification: 126 × 87 × 113 mm, 1 L) was used to construct the light-proof culture system, and 96-well plates were used for planting. The Hoagland’s nutrient solution was adopted for growing rice in hydroponics. A 1/2 concentration nutrient solution was used for the first 3 d after planting, then replaced with full concentration from the 4th day, and the liquid level maintained at a 2/3 submerged state of the root system. The evaporative water loss was replenished every 48 hours, the nutrient solution was renewed every 72 hours, and the pH was adjusted to 5.8±0.2. The treatment group was inoculated with 3% sterilized MAA fermentation supernatant, and the control group was supplemented with the same amount of sterile PDB medium. Three biological replicates of each treatment were established under a photoperiod of 16 h/8 h (day/night) and a temperature of 28 °C. The shoot height of rice plants was measured after 7 d, and the root systems were gently rinsed and straightened before measuring the length of the longest root using a straightedge at 7 days post-treatment as the root length.

### MAA Colonization Confirmation

To assess the extent of the endophytic association (MAA), root samples from the treatment and control group after 7 d were collected. DNA extraction and PCR amplification of the ITS region were subsequently performed. The relative abundance of fungal communities was assessed using high-throughput sequencing of the ITS region, which was amplified using the primers ITS1 and ITS4, and PCR products were sequenced on an Illumina MiSeq platform. Raw sequences were processed using the QIIME2 pipeline, including quality filtering, denoising, and clustering into ASVs. Taxonomic assignment was performed using the UNITE database. Relative abundances were calculated as the proportion of each ASV relative to the total number of sequences per sample. Statistical analyses, including PERMANOVA and PCoA, were conducted to compare fungal communities across different treatments.

### Effects of different culture media on *M. robertsii* growth and Rice growth promotion

Under the basic culture conditions of carbon source (glucose), nitrogen source (peptone), pH (6.0), cultivation time (7 d), β-Alanine (β-Ala, 0.2%) and Al_2_(SO_4_)_3_ (50 μmol/L), single-factor experiments were conducted by altering one variable at a time. The tested parameters included: carbon sources (glucose, sucrose, galactose, corn starch, ammonium sulfate, and soluble starch) to replace glucose in the basic medium; nitrogen sources (yeast extract, L-glutamic acid, beef extract, KNO_3_, and peptone) to replace peptone in the basic medium; pH (5.0, 6.0, 7.0, 8.0, and 9.0); cultivation time (7 d, 9 d, 11 d, 13 d, and 15 d); β-Ala concentration (0.1%, 0.2%, 0.3%, 0.4%, and 0.5%); and Al_2_(SO_4_)_3_ concentration (10 μmol/L, 20 μmol/L, 50 μmol/L, 100 μmol/L, and 150 μmol/L). The composition of different culture media is shown in Table 1. All treatments were performed in triplicate and incubated in a rotary shaker at 220 rpm for 7 days.

**Table 1.** Effects of Six factors and five horizontal medium factors on the growth of MAA and Rice.

| Factor | Carbon Source | Nitrogen Source | Fermentation Time ( d ) | β-Ala Concentration (%) | Al<sub>2</sub>(SO<sub>4</sub>)<sub>3</sub> Supplementation (μmol/L) | pH |
| --- | --- | --- | --- | --- | --- | --- |
| Levels |  |  |  |  |  |  |
| 1 | glucose | peptone | 7 | 0.1 | 10 | 5.0 |
| 2 | soluble starch | yeast | 9 | 0.2 | 20 | 6.0 |
| 3 | corn flour | glutamic acid | 11 | 0.3 | 50 | 7.0 |
| 4 | galactose | (NH<sub>4</sub>)<sub>2</sub>SO<sub>4</sub> | 13 | 0.4 | 100 | 8.0 |
| 5 | sucrose | KNO<sub>3</sub> | 15 | 0.5 | 150 | 9.0 |

Fungal growth assessment: To evaluate the impact of different culture media on fungal growth, solid media were prepared and the components are shown in Table 1. Each solid medium was poured into sterile Petri dishes and allowed to solidify. After solidification in Petri dishes, 2 µL of conidial suspension (1 × 10^7^ CFU/mL) was centrally inoculated and incubated at 28°C in darkness. Each treatment was performed in triplicate. Fungal growth was assessed by measuring the colony diameter after 7 days. Additionally, the morphology of the colonies (e.g., color, texture, and sporulation) was recorded.

Rice growth promotion assay: The effects of MAA cultured with different media (Table 1) on rice growth were evaluated hydroponically using Hoagland nutrient solution. Each treatment included 96 rice seedlings. The shoot height of rice plants was measured after 7 d, and the root systems were gently rinsed and straightened before measuring the length of the longest root using a straightedge at 7 days post-treatment as the root length.

### Single-Factor Test of the Effects of Different Fermentation Conditions on the Growth Promoting Activity of MAA

Optimization was carried out through single-factor experiments: key variables including carbon source, nitrogen source, pH, cultivation time, β-Ala concentration and Al_2_(SO_4_)_3_ supplementation were individually tested under basic conditions (as described in the previous section). A total of 36 distinct medium formulations were designed to systematically evaluate each factor, with detailed compositions provided in Appendix Table S4. The growth-promoting activity was evaluated by comparing the plant height of 7-day-old rice seedlings and the plant height and root length of 15-day-old seedlings.

### Response Surface Optimization Test

Based on the single-factor test results, key parameters including sucrose concentration (A), pH (B), β-Ala concentration (C), and Al_2_(SO_4_)_3_ supplementation (D) were further optimized. The response surface methodology (RSM) was employed following established protocols (23, 34), with the experimental design detailed in Tables S1 and S2. The optimization results were evaluated based on the 7-day root length of rice.

### Evaluation of Rice Growth-Promoting Traits

Additional growth-promoting characteristics were assessed, including chlorophyll content, adventitious root number and root hair phenotype.

#### Chlorophyll content

The relative chlorophyll content was determined using a SPAD-502 Plus chlorophyll meter, with measurements taken at a standardized position on the leaves.

#### Adventitious root number

After 15 days of hydroponic cultivation, the adventitious roots were counted for each plant (n = 96). The adventitious root number was calculated as the average number of adventitious roots distributed along the crown root.

#### Root hair phenotype

To quantify root hair development, 7-day-old roots were imaged in the root maturation zone using an FV1000 confocal microscope (Olympus).

The growth rate of each parameter was calculated as: growth rate (%) = [(treatment group value - control group value) / control group value] × 100.

## Statistical Analysis

All experiments were performed with at least three biological replicates. Data are presented as means ±standard error (SE), and Origin 2022b was used for mapping (Origin Lab, Northampton, MA, USA). Statistical significance among treatments was determined by one-way ANOVA followed by Fisher’s Least Significant Difference (LSD) post-hoc test (P < 0.05) using SPSS 22.0 (IBM SPSS Statistics, Chicago, IL, USA). Significant differences between treatment groups are indicated by different superscript letters.

## References

1. Hu S, Bidochka MJ. 2021. Root colonization by endophytic insect-pathogenic fungi. Journal of Applied Microbiology 130:570–581.

2. Hu G, Leger RJS. 2002. Field Studies Using a Recombinant Mycoinsecticide (Metarhizium anisopliae) Reveal that It Is Rhizosphere Competent. Applied and Environmental Microbiology 68:6383–6387.

3. Wakil W, Boukouvala MC, Kavallieratos NG, Naeem A, Ntinokas D, Ghazanfar MU, Avery PB. 2025. The Inevitable Fate of Tetranychus urticae on Tomato Plants Treated with Entomopathogenic Fungi and Spinosad. Journal of Fungi 11:138.

4. Menino GC, Tanaka FA, Zambrosi FC. 2025. Preventive foliar application of sparingly soluble copper source is effective to protect against Asian soybean rust. Australasian Plant Pathology 54:25–31.

5. Ahmad I, Jimenez-Gasco MdM, Barbercheck ME. 2024. Water stress and black cutworm feeding modulate plant response in maize colonized by Metarhizium robertsii. Pathogens 13:544.

6. Wakil W, Kavallieratos NG, Eleftheriadou N, Haider SA, Qayyum MA, Tahir M, Rasool KG, Husain M, Aldawood AS. 2024. A winning formula: Sustainable control of three stored-product insects through paired combinations of entomopathogenic fungus, diatomaceous earth, and lambda-cyhalothrin. Environmental Science and Pollution Research 31:15364–15378.

7. Liao X, O’Brien TR, Fang W, St. Leger RJ. 2014. The plant beneficial effects of Metarhizium species correlate with their association with roots. Applied microbiology and biotechnology 98:7089–7096.

8. Jaber LR, Araj S-E. 2018. Interactions among endophytic fungal entomopathogens (Ascomycota: Hypocreales), the green peach aphid Myzus persicae Sulzer (Homoptera: Aphididae), and the aphid endoparasitoid Aphidius colemani Viereck (Hymenoptera: Braconidae). Biological Control 116:53–61.

9. Ahmad I, del Mar Jiménez-Gasco M, Luthe DS, Shakeel SN, Barbercheck ME. 2020. Endophytic Metarhizium robertsii promotes maize growth, suppresses insect growth, and alters plant defense gene expression. Biological Control 144:104167.

10. Behie S, Zelisko P, Bidochka M. 2012. Endophytic insect-parasitic fungi translocate nitrogen directly from insects to plants. Science 336:1576–1577.

11. Sasan RK, Bidochka MJ. 2013. Antagonism of the endophytic insect pathogenic fungus Metarhizium robertsii against the bean plant pathogen Fusarium solani f. sp. phaseoli. Canadian journal of plant pathology 35:288–293.

12. Chairin T, Petcharat V. 2017. Induction of defense responses in longkong fruit (Aglaia dookkoo Griff.) against fruit rot fungi by Metarhizium guizhouense. Biological Control 111:40–44.

13. Elena GJ, Beatriz PJ, Alejandro P, Lecuona R. 2011. Metarhizium anisopliae (Metschnikoff) Sorokin promotes growth and has endophytic activity in tomato plants. Adv Biol Res 5:22–27.

14. Jiang X, Fang W, Tong J, Liu S, Wu H, Shi J. 2022. Metarhizium robertsii as a promising microbial agent for rice in situ cadmium reduction and plant growth promotion. Chemosphere 305:135427.

15. Jiang X, Dai J, Zhang X, Wu H, Tong J, Shi J, Fang W. 2022. Enhanced Cd efflux capacity and physiological stress resistance: The beneficial modulations of Metarhizium robertsii on plants under cadmium stress. Journal of Hazardous Materials 437:129429.

16. Chowdhury MZH, Mostofa MG, Mim MF, Haque MA, Karim MA, Sultana R, Rohman MM, Bhuiyan A-U-A, Rupok MRB, Islam SMN. 2024. The fungal endophyte Metarhizium anisopliae (MetA1) coordinates salt tolerance mechanisms of rice to enhance growth and yield. Plant physiology and biochemistry 207:108328.

17. Luo F, Tang G, Hong S, Gong T, Xin X-F, Wang C. 2023. Promotion of Arabidopsis immune responses by a rhizosphere fungus via supply of pipecolic acid to plants and selective augment of phytoalexins. Science China Life Sciences 66:1119–1133.

18. Barelli L, Behie SW, Hu S, Bidochka MJ. 2022. Profiling destruxin synthesis by specialist and generalist Metarhizium insect pathogens during coculture with plants. Applied and Environmental Microbiology 88:e02474–21.

19. Kumar CS, D’Silva S, Praveena R, Kaprakkaden A, Krishnan LA, Rajkumar MB, Srinivasan V, Dinesh R. 2024. Zinc solubilization and organic acid production by the entomopathogenic fungus, Metarhizium pingshaense sheds light on its key ecological role in the environment. Science of the Total Environment 923:171348.

20. Conrado R, Gomes TC, Roque GSC, De Souza AO. 2022. Overview of bioactive fungal secondary metabolites: cytotoxic and antimicrobial compounds. Antibiotics 11:1604.

21. Singh V, Haque S, Niwas R, Srivastava A, Pasupuleti M, Tripathi C. 2017. Strategies for fermentation medium optimization: an in-depth review. Frontiers in microbiology 7:2087.

22. Wang YH, Feng JT, Zhang Q, Zhang X. 2008. Optimization of fermentation condition for antibiotic production by Xenorhabdus nematophila with response surface methodology. Journal of Applied Microbiology 104:735–744.

23. He J, Zhang X, Wang Q, Li N, Ding D, Wang B. 2023. Optimization of the fermentation conditions of Metarhizium robertsii and its biological control of wolfberry root rot disease. Microorganisms 11:2380.

24. Melini F, Luziatelli F, Bonini P, Ficca AG, Melini V, Ruzzi M. 2023. Optimization of the growth conditions through response surface methodology and metabolomics for maximizing the auxin production by Pantoea agglomerans C1. Frontiers in Microbiology 14:1022248.

25. Martín JF, Liras P. 2024. Diamine Fungal Inducers of Secondary Metabolism: 1,3-Diaminopropane and Spermidine Trigger Enzymes Involved in β-Alanine and Pantothenic Acid Biosynthesis, Precursors of Phosphopantetheine in the Activation of Multidomain Enzymes. Antibiotics 13:826.

26. Kohli PS, Maurya K, Thakur JK, Bhosale R, Giri J. 2022. Significance of root hairs in developing stress-resilient plants for sustainable crop production. Plant, Cell & Environment 45:677–694.

27. Ramesh K, Chandrasekaran B, Balasubramanian T, Bangarusamy U, Sivasamy R, Sankaran N. 2002. Chlorophyll dynamics in rice (Oryza sativa) before and after flowering based on SPAD (chlorophyll) meter monitoring and its relation with grain yield. Journal of Agronomy and Crop Science 188:102–105.

28. Zhou W, Fan L, Gao S, Zhou S, Shi G. 2025. Enhancing Seedling Growth and Root Development Through Symbiotic Interactions: The Role of Metarhizium anisopliae in Tree Peony Root Systems. Journal of Plant Growth Regulation:1–14.

29. da Silva PF, dos Santos MSN, Araújo BdA, Kerber BD, de Oliveira HAP, Guedes JVC, Mazutti MA, Tres MV, Zabot GL. 2025. Co-Cultivations of Beauveria Bassiana, Metarhizium Anisopliae, and Trichoderma Harzianum to Produce Bioactive Compounds for Application in Agriculture. Fermentation 11:30.

30. Cao W, Yu C, Zhao Y, Lin Q, Deng C, Li C. 2024. Biological characteristics, artificial domestication conditions optimization, and bioactive components of Beauveria caledonica. Microorganisms 12:1554.

31. Liu Y, Li J, Shi J, Pan Y, Yang S, Xue Y. 2024. Combined metabolome and transcriptome analysis reveals the key pathways involved in the responses of soybean plants to high Se stress. Ecotoxicology and Environmental Safety 287:117262.

32. Chen J, Ullah C, Giddings Vassão D, Reichelt M, Gershenzon J, Hammerbacher A. 2021. Sclerotinia sclerotiorum infection triggers changes in primary and secondary metabolism in Arabidopsis thaliana. Phytopathology® 111:559–569.

33. Kaur S, Campbell BJ, Suseela V. 2022. Root metabolome of plant–arbuscular mycorrhizal symbiosis mirrors the mutualistic or parasitic mycorrhizal phenotype. New Phytologist 234:672–687.

34. Gao X, He Q, Jiang Y, Huang L. 2016. Optimization of nutrient and fermentation parameters for antifungal activity by Streptomyces lavendulae Xjy and its biocontrol efficacies against Fulvia fulva and Botryosphaeria dothidea. Journal of Phytopathology 164:155–165.

